# A machine learning tool for interpreting differences in cognition using brain features

**DOI:** 10.1101/558403

**Authors:** Tiago Azevedo, Luca Passamonti, Pietro Lió, Nicola Toschi

## Abstract

Predicting variability in cognition traits is an attractive and challenging area of research, where different approaches and datasets have been implemented with mixed results. Some powerful Machine Learning algorithms employed before are difficult to interpret, while other algorithms are easy to interpret but might not be as powerful. To improve understanding of individual cognitive differences in humans, we make use of the most recent developments in Machine Learning in which powerful prediction models can be interpreted with confidence. We used neuroimaging data and a variety of behavioural, cognitive, affective and health measures from 905 people obtained from the Human Connectome Project (HCP). As a main contribution of this paper, we show how one could interpret the neuroanatomical basis of cognition, with recent methods which we believe are not yet fully explored in the field. By reducing neuroimages to a well characterised set of features generated from surface-based morphometry and cortical myelin estimates, we make the interpretation of such models easier as each feature is self-explanatory. The code used in this tool is available in a public repository: https://github.com/tjiagoM/interpreting-cognition-paper-2019

## 1 Introduction

Predicting variability in cognitive functioning can have important consequences in terms of delineating the life trajectory of a person. Individual differences in cognition have been related to important outcome measures like education or occupational achievement, general health, longevity, and risk to develop dementia. One way of predicting cognitive performances at the single subject level is to use different sources of neuroimaging data that assess different aspects of brain function and structure (anatomy). Resting-state functional connectivity [11][3] or neuro-anatomically extracted features [13] are examples of such sources of neuroimaging data.

However, no one in this field so far has tried to capitalise on multi-modal neuroimaging, i.e. on the possibility to predict individual differences in cognition by simultaneously using different type of information regarding the brain structure or function. In this study, we combined different structural neuroimaging modalities (T1-weighted and T1/T2-derived intracortical myelin estimates) to predict subject-specific scores in a series of cognitive and demographic measures derived from a data-reduction analyses of a large set of behavioural measures, drawn from the Human Connectome Project (HCP).

Machine Learning has recently gained attention in a number of applied biomedical disciplines due to the increased prediction power it can provide. In specifically, neural networks and deep learning methods are the most well-known but they usually bring a lot of complexity when one wants to interpret the results or avoid overfitting [15]. Instead, we focused our attention in a type of gradient boosting decision tree algorithm, XGBoost [1], which have recently achieved top results in applied machine learning competitions using tabular data. As it is built on top of decision trees, it also has the advantage that there is space to try to understand how the model makes its decisions.

To interpret the neuroanatomical basis of cognitive measures, we use recently developed algorithms that improve the interpretability capacity of machine learning models without neglecting their prediction power. In specific, we used an adaptation of SHAP (SHapley Additive exPlanations) [9] which is a unified framework for interpreting predictions without falling in common mistakes that do not bring consistent and true interpretations [10] .

Interpretation of 3D neuroimages can be complex, and usually relies on visual plots and specific neuroanatomical knowledge for interpretation. In this paper, we reduced the different structural neuroimaging modalities by using surface-based morphometry and parcellated cortical myelin estimates as subject-wise features in XGBoost. By reducing multi-modal neuroimaging measures to a well characterised set of features, we make the interpretation of such models easier as each feature is self-explanatory *per se*. For example, when interpreting machine learning models that use 3D images as a basis, it is possible to visually identify areas of the brain responsible for the prediction, but it is not obvious whether those regions are related, for instance, to actual values of thickness or surface area.

The main contribution of this paper is thus to show how one could interpret the neuroanatomical basis of cognition, by applying state-of-the-art machine learning methods which we believe are not yet fully and correctly explored in the field. The code used in the resulting tool is publicly available: https://github.com/tjiagoM/interpreting-cognition-paper-2019

## 2 Methods

### 2.1 Dataset and Participants

Pre-processed structural Magnetic Resonance Images as well as demographic and cognitive data from 1200 subjects were obtained from the Human Connectome Project (HCP) public repository^5^.

After accounting for missing information, the total number of subjects included in our analysis was 905, where around 53% were females, and 47% were males. All participants were young and healthy adults, with a median age of 29, with no hypertension, alcohol misuse, panic disorder, depression, or other psychiatric and neurologic disorder, or history of childhood conduct problems. The majority of people were right-handed white americans with a non-hispanic or latino background (cf. Table 1).

**Table 1:** Demographic Information of the 905 people analysed

|  | 1<sup>st</sup><br>Quantile | Median | 3<sup>rd</sup><br>Quantile |
| --- | --- | --- | --- |
| Age (in years) | 26 | 29 | 32 |
| Weight (in kg) | 63.96 | 75.30 | 88.56 |
| Height (in metres) | 1.651 | 1.702 | 1.778 |
| Years of education completed | 14 | 16 | 16 |
| Body Mass Index | 22.715 | 25.375 | 29.135 |
| Systolic Blood Pressure (in mmHg) | 114 | 123 | 132 |
| Diastolic Blood Pressure (in mmHg) | 70 | 76 | 83 |
| Number of childhood conduct problems | 0 | 0 | 1 |
| Number of panic disorder symptoms | 0 | 0 | 0 |
| Number of depressive symptoms | 0 | 0 | 0 |
| Total drinks in the past 7 days | 0 | 2 | 6.5 |
| **Gender** | Female: 53%<br>Male: 47% |  |  |
| **Race** | American Indian/Alaskan Natural: 0.2%<br>Asian/Natural Hawaiian/Other Pacific Island: 5.7%<br>Black or African American: 14.6%<br>More than one: 2.4%<br>Unknown or Not Reported: 1.5%<br>White: 75.5% |  |  |
| **Ethnicity** | Hispanic/Latino: 9.1%<br>Not Hispanic/Latino: 89.8%<br>Unknown or Not Reported: 1.1% |  |  |
| **Handedness** | Right-handed: 90.8%<br>Left-handed: 9.0%<br>Mixed: 0.2% |  |  |

### 2.2 Data and Image Processing

HCP structural T1w images were collected from a 3-Tesla Siemens Skyra unit (housed at Washington University in St. Louis) using an axial T1-weighted sequence (TR = 2400 ms, TE = 2.14 ms, flip angle=8, voxel-size 0.7 0.7 0.7 mm3). The T1w data was passed through the full Freesurfer (v. 5.3)^6^ reconstruction stream to calculate cortical thickness, surface area, number of vertices in the cortex, gray volume, integrated rectified mean curvature, integrated rectified gaussian curvature, folding index, and intrinsic curvature index. The optimal pipeline used to obtain these segmentation is described in detail in a previous article [6]. To map all subjects’ brains to a common space, namely the Desikan-Killiany Atlas [2], reconstructed surfaces were registered to an average cortical surface atlas released by HCP^7^ using a non-linear procedure that optimally aligned sulcal and gyral features across subjects [4].

Myelin maps were generated by the HCP consortium according to Glasser and Van Essen (2011) [5]. Transformation of myelin map data from the individual subject’s native mesh to the right fsaverage template (LR) standard mesh involves two deformation maps, one representing registration from the native mesh to the fsaverage left mesh (L) and fsaverage right mesh (R) and another representing registration between L and R and LR. The two deformation maps were concatenated into a single deformation map using Caret software that was applied to the individual subject’s myelin map data, cortical thickness data, and surface curvature data. The individual myelin map data were normalised to a group global mean and then averaged at each surface node in order to achieve anatomical correspondence to other structural features.

### 2.3 Factor Analysis

We extracted 9 factors which explained 70% of cumulative variance using Principal Component Analysis (PCA). Hence, twenty-three measures of affect, cognition, and health including self-report questionnaires, neuropsychological tasks, and other behavioural indices from the HCP database were grouped into independent and latent components of emotional behaviour, affect, psychological well-being, cognitive functioning, and general health using factor analysis (FA). Subject-specific loading values were employed as dependent variables. In Table 2 it is possible to see what each factor is representing.

**Table 2:** Loading values for n=23 behavioural variables on the estimated factors from the factor analysis. The colour scale represents the item loading scores on the independent factors

|  | Fluid Intelligence | Visual Memory | Sex-related factor | Executive Functions | Sustained Attention | Waiting Impulsivity | Linguistic Skills | Verbal Episodic Memory | Visuo-spatial Skills |
| --- | --- | --- | --- | --- | --- | --- | --- | --- | --- |
| Total Intracranial Volume | 0.121 | 0.138 | -0.772 | 0.107 | 0.042 | 0.071 | -0.033 | -0.005 | 0.093 |
| Gender | -0.048 | 0.047 | 0.841 | -0.041 | -0.058 | -0.010 | 0.005 | -0.082 | -0.001 |
| Age | -0.084 | 0.182 | 0.455 | 0.077 | 0.137 | 0.067 | -0.019 | 0.471 | 0.058 |
| Memory - Picture Sequence Memory Test Score | 0.124 | 0.560 | 0.220 | 0.130 | 0.036 | -0.034 | -0.035 | -0.269 | 0.215 |
| Executive Function/Cognitive Flexibility - Card Sort Test Score | 0.102 | 0.187 | 0.001 | 0.790 | 0.058 | -0.025 | -0.029 | -0.008 | -0.006 |
| Executive Function/Inhibition - Flanker Inhibitory Control and Attention Test Score | -0.016 | 0.041 | -0.156 | 0.800 | 0.005 | 0.025 | -0.109 | -0.011 | 0.011 |
| Fluid Intelligence - Penn Progressive Matrices: Number of Correct Responses | 0.898 | 0.272 | -0.117 | 0.110 | 0.030 | 0.083 | -0.105 | -0.127 | -0.010 |
| Penn Progressive Matrices: Total Skipped Items | -0.901 | -0.260 | 0.111 | -0.106 | -0.021 | -0.065 | 0.088 | 0.109 | 0.007 |
| Penn Progressive Matrices: Median Reaction Time for Correct Responses | 0.875 | 0.071 | -0.050 | -0.096 | 0.028 | 0.118 | -0.017 | 0.103 | -0.003 |
| Oral Reading Recognition Test Score | -0.070 | -0.083 | -0.028 | -0.013 | 0.018 | 0.002 | 0.865 | 0.016 | 0.035 |
| Picture Vocabulary Test Score | -0.076 | -0.003 | 0.061 | -0.018 | -0.023 | -0.027 | 0.871 | 0.016 | -0.010 |
| Processing Speed Test Score | -0.005 | 0.101 | 0.021 | 0.732 | 0.060 | 0.018 | 0.103 | -0.182 | -0.039 |
| Delay Discounting: Area Under the Curve for Discounting of \$200 | 0.101 | 0.047 | -0.035 | -0.005 | 0.015 | 0.900 | -0.038 | -0.059 | 0.036 |
| Delay Discounting: Area Under the Curve for Discounting of \$40,000 | 0.111 | 0.102 | -0.037 | 0.021 | 0.006 | 0.898 | 0.011 | -0.023 | -0.010 |
| Spatial Orientation - Variable Short Penn Line Orientation: Total Number Correct | 0.215 | 0.665 | -0.413 | 0.094 | 0.062 | 0.102 | -0.042 | -0.013 | -0.182 |
| Variable Short Penn Line Orientation: Median Reaction Time Divided by Expected Number of Clicks for Correct | -0.017 | 0.013 | -0.050 | -0.034 | -0.013 | 0.029 | 0.027 | 0.042 | 0.940 |
| Variable Short Penn Line Orientation: Total Positions Off for All Trials | -0.242 | -0.668 | 0.420 | -0.092 | -0.061 | -0.125 | 0.029 | 0.008 | 0.189 |
| Sustained Attention - Short Penn Continuous Performance Test: Sensitivity | 0.065 | 0.152 | -0.021 | 0.095 | 0.905 | 0.027 | 0.005 | -0.027 | -0.017 |
| Short Penn Continuous Performance Test: Specificity | 0.113 | 0.566 | 0.248 | 0.066 | -0.064 | 0.048 | 0.067 | 0.155 | -0.039 |
| Short Penn Continuous Performance Test: Longest Run of Non-Responses | 0.000 | 0.101 | 0.053 | -0.025 | -0.923 | 0.006 | 0.009 | 0.063 | -0.003 |
| Verbal Episodic Memory - Penn Word Memory Test: Total Number of Correct Responses | 0.085 | 0.252 | 0.148 | -0.007 | 0.158 | 0.073 | -0.029 | -0.643 | -0.019 |
| Penn Word Memory Test: Median Reaction Time for Correct Responses | 0.033 | 0.006 | 0.036 | -0.228 | -0.007 | -0.049 | 0.015 | 0.761 | 0.008 |
| List Sorting Working Memory Test Score | 0.115 | 0.586 | -0.066 | 0.124 | 0.028 | 0.043 | -0.104 | -0.132 | 0.112 |

### 2.4 Cross-Validation Procedure

To predict each factor using our generated features, we implemented a Nested Cross-Validation (CV) procedure as depicted in Figure 1, with 5 outer folds, and 3 inner folds. Essentially, the data is divided in 5 folds, and selecting each fold once, the other four folds are selected to run an inner loop. For each inner loop the data is divided in 3 folds, where 2 of these folds are used for feature selection and hyperparameter search, selecting the hyperparameters that yield the best Mean Squared Error (MSE) in the remaining fold. After running this process for each one of the 3 folds, the model that yield the lowest mean MSE is selected and used in the fold selected in that outer loop. In other words, this process selects 5 different models for each outer fold.

**Fig. 1:**
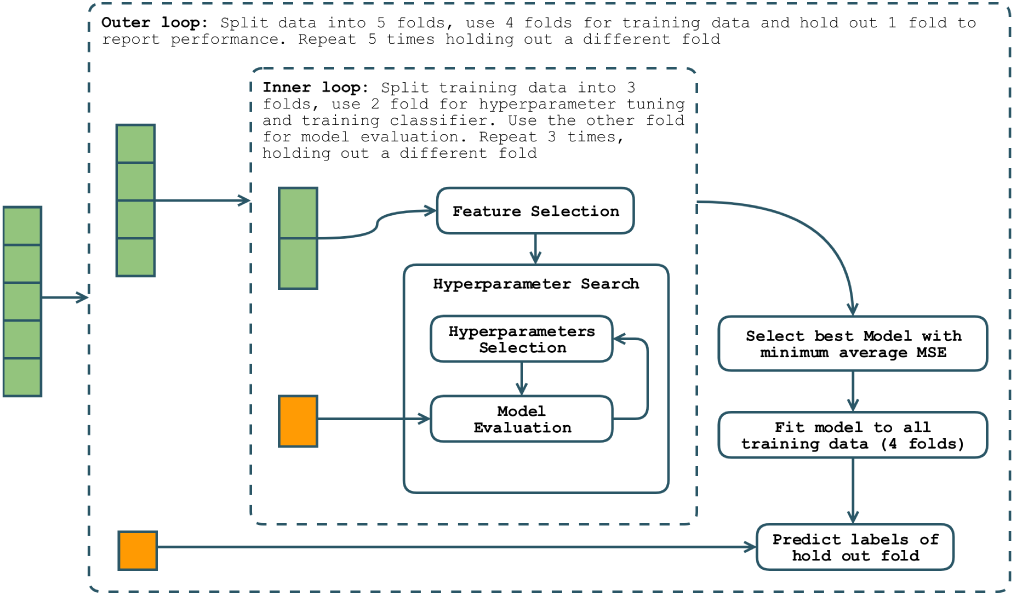
Illustration of the Nested Cross-Validation used in this paper. The coloured boxes represent the data

We use XGBoost [1] as our machine learning method to predict our factors. XGBoost is a effective, scalable gradient boosting tree algorithm, used worldwide in machine learning and data science competitions to achieve the best predicting results in tabular data. It uses several decision trees (models) as an ensemble added together to make the final prediction. It is called gradient boosting because it uses a gradient descent algorithm to minimise the loss when a new tree/method is added.

Regarding the Feature Selection step, in each iteration we selected the 450 values that share more mutual information with the factor being predicted. We used the scikit-learn [12] implementation as originally described in Kraskov et al. [8] and Ross’ [14] papers.

### 2.5 Feature Importance Analysis

Given the complexity of the XGBoost, and the reported issues when understanding how the many decision trees are used for prediction, we used the recent SHAP [10] (SHapley Additive exPlanations), a unified framework for interpreting predictions. Specifically, we used the recent adaptation for tree ensemble methods [9]. SHAP solves reports that popular feature attribution methods are inconsistent and incapable of reporting the true impact of features in tree ensemble methods like XGBoost.

## 3 Results

Table 3 summarises the results of the nested CV procedure over the 9 factors, by reporting the averaged Mean Squared Errors (MSE) and Pearson-r correlations between actual and predicted values, with the respective standard deviations. In this Section we will only report results on Factors 1, 2, 3 and 7 as the remaining factors either had a high MSE standard deviation or Pearson-r correlation too close to zero.

**Table 3:** Prediction Power regarding Mean Squared Error and Pearson-r correlation. In bold the only factors that are further analysed in this paper

| Factor | Mean MSE<br>(std deviation) | Mean Pearson-r<br>(std deviation) |
| --- | --- | --- |
| **Factor 1 - Fluid Intelligence** | 0.932 (0.063) | 0.138 (0.024) |
| **Factor 2 - Visual Memory** | 0.924 (0.071) | 0.114 (0.031) |
| **Factor 3 - Sex-related Factor** | 0.407 (0.038) | 0.763 (0.022) |
| Factor 4 - Executive Functions | 0.961 (0.115) | 0.171 (0.072) |
| Factor 5 - Sustained Attention | 1.072 (0.873) | -0.038 (0.044) |
| Factor 6 - Waiting Impulsivity | 1.115 (0.305) | 0.142 (0.086) |
| **Factor 7 - Linguistic Skills** | 0.941 (0.055) | 0.119 (0.056) |
| Factor 8 - Verbal Episodic Memory | 1.015 (0.079) | 0.066 (0.041) |
| Factor 9 - Visuo-spatial Skills | 0.988 (0.101) | -0.031 (0.048) |

Figure 2 summarises another powerful interpretation that one can get from the SHAP method when applied to a prediction model. Each feature is assigned a SHAP value which represents both the magnitude and direction of its contribution. Its absolute value indicates how powerful that feature is to the prediction, and its signal expresses whether it contributes towards a higher or lower predicted value.

**Fig. 2:**
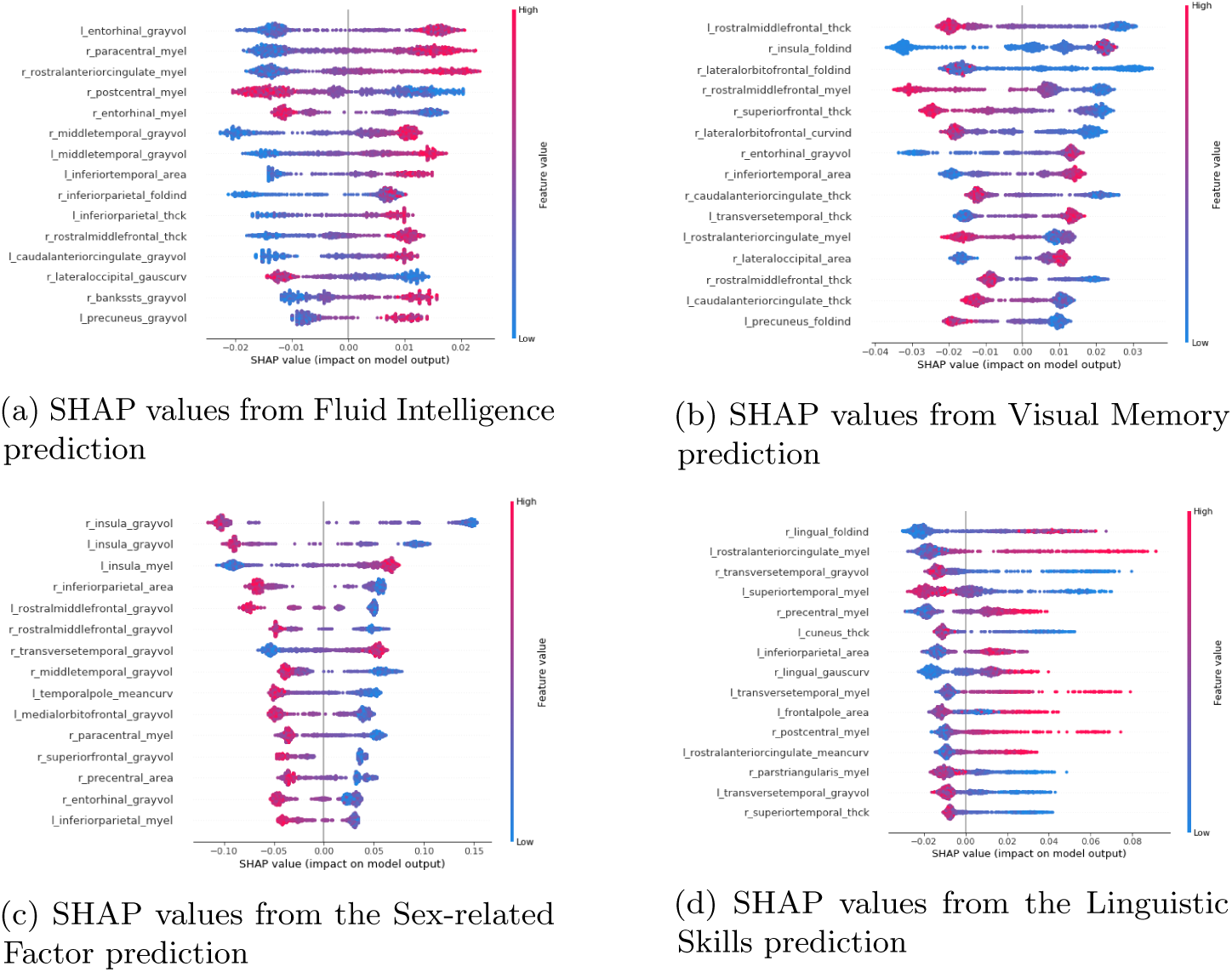
Contribution of the most important features in each factor, from one of the respective outer folds. For each feature, vertical dispersion stands for the data points who share the same SHAP value for that feature. Higher SHAP values mean they contribute in a positive direction to the final predicted variable

The median rank of brain features when predicting Fluid Intelligence is not very consistent across the outer folds, as it is possible to see in Table 4 where most of the features identified were ranked, on average, below 10, and the Median Absolute Deviation is quite high as well. Some of the brain areas identified are situated around the temporal lobes (middle, inferior and entorhinal/medial, as well as parahippocampal cortex), in which surface areas and gray volumes were more important to predict Fluid Intelligence. Some other areas around the brain were selected, representing not only gray volumes but also folding indexes and myelin values. The left hemisphere is selected much more times than the right hemisphere.

**Table 4:**
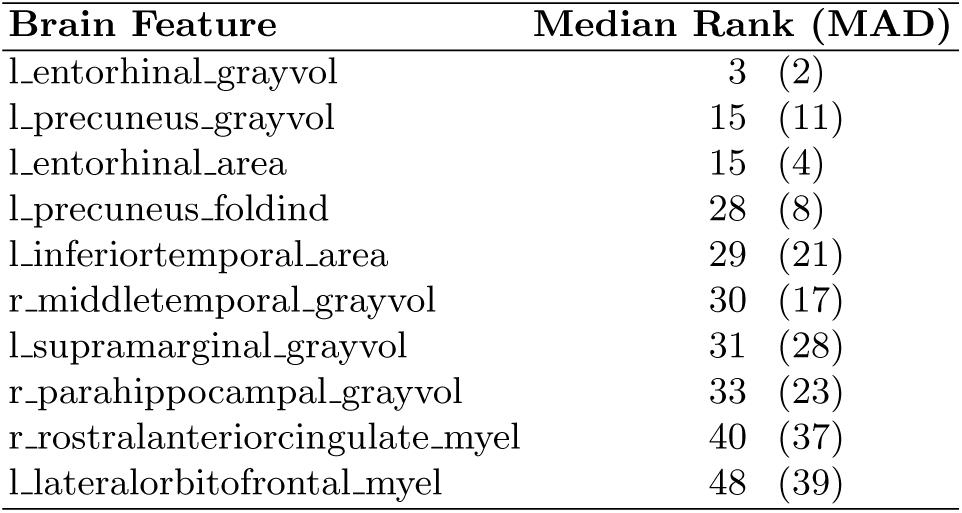
Median rank of the ten most important features when predicting Fluid Intelligence. We considered features that appeared at least in 4 folds. MAD stands for Median Absolute Deviation

As one can see from the most important features in one outer fold when predicting Fluid Intelligence (cf. Figure 2a), generally a higher value of that feature contributes to a higher value of Fluid Intelligence, with a clear exception of the myelin of the Post-Central and Entorhinal area.

The most important features selected across the five outer folds for the prediction of Visual Memory are much more consistent across the outer folds, as the median ranks of the 10 most important features are below 20 (cf. Table 5) and generally the Median Absolute Deviation is below 10. For this factor the most important features are clearly the folding index of the insula and the gray volume of the entorhinal cortex. Frontal lobes play an important role, by having features being selected for the superior frontal, caudal middle frontal, rostral middle frontal, lateral orbitofrontal, and pars opercularis cortex regions, mostly with cortical thicknesses selected. In contrast with Fluid Intelligence, the right hemisphere is selected much more times to predict Visual Memory, which is consistent with previous findings linking the right hemisphere to visual memory [7], although the gray volumes of the entorhinal region were selected in both hemispheres.

**Table 5:** Median rank of the ten most important features when predicting Visual Memory. We considered features that appeared at least in 4 folds. MAD stands for Median Absolute Deviation

| **Brain Feature** | **Median Rank (MAD)** |
| --- | --- |
| r_insula_foldind | 2 (1) |
| l_entorhinal_grayvol | 3 (2) |
| r_superiorfrontal_thck | 7 (2) |
| r_entorhinal_grayvol | 7 (6) |
| r_caudalmiddlefrontal_thck | 7 (1) |
| r_cuneus_thck | 9 (5) |
| l_rostralmiddlefrontal_thck | 12 (7) |
| r_lateralorbitofrontal_foldind | 16 (13) |
| r_parsopercularis_thck | 16 (7) |
| r_inferiortemporal_area | 20 (18) |

As one can see from Figure 2b, when most of these features have a low value, the predicted Visual Memory will be higher. There are some exceptions, but an highlight is the behaviour of the most important feature in terms of median rank (Insula’s folding index), in which lower values might increase or decrease Visual Memory.

Prediction of the sex-related factor yielded not only the best pearson-r correlation values, but the most consistent median ranks across the outer folds, with the 10 most important features always being in top 15 (cf. Table 6). Insula plays a big role in distinguishing a male-or female-like brain, specifically with regards to its gray volume in both hemispheres, as well as the myelin in the left hemisphere. After the insula, the frontal lobes are more prominent, with the presence of features extracted from the medial orbitofrontal cortex, pre-central and rostral middle frontal areas. There is a clear dominance of the left hemisphere in predicting this factor.

**Table 6:** Median rank of the ten most important features when predicting the Sex-related factor. We considered features that appeared at least in 4 folds. MAD stands for Median Absolute Deviation

| **Brain Feature** | **Median Rank (MAD)** |
| --- | --- |
| r_insula_grayvol | 1 (0) |
| l_insula_grayvol | 3 (1) |
| l_medialorbitofrontal_grayvol | 3 (2) |
| r_inferiorparietal_area | 4 (1) |
| l_temporalpole_meancurv | 8 (1) |
| l_insula_myel | 8 (5) |
| l_transversetemporal_myel | 11 (2) |
| l_precentral_myel | 12 (5) |
| l_rostralmiddlefrontal_grayvol | 13 (8) |
| r_precentral_area | 13 (11) |

From Figure 2c it is possible to find a clear trend in which a lower feature value help in a lower Sex-related factor or, in other words, to a brain looking more like a female one. It is also possible to see that the gray volumes of the insula in both hemispheres have a higher SHAP value, in magnitude, than any other feature which, adding to its median rank, shows how important that feature is in distinguishing male-and female-like brains.

Finally, the median ranks of the 10 most important features when predicting Linguistic Skills are not very consistent across the outer folds, but not as inconsistent as when predicting Fluid Intelligence (cf. Table 7). The presence of regions from the frontal lobes is significant, namely the pre-central, rostral anterior cingulate, frontal pole, and caudal anterior cingulate regions. However, the most significant regions are around the temporal lobes: superior temporal, transverse temporal, lingual, parahippocampal, and middle temporal. As opposed to the other factors analysed, there is no clear dominance of one hemisphere over the other.

**Table 7:** Median rank of the ten most important features when predicting Linguistic Skills. We considered features that appeared at least in 4 folds. MAD stands for Median Absolute Deviation

| Brain Feature | Median Rank (MAD) |
| --- | --- |
| r_precentral_myel | 8 (4) |
| l_superiortemporal_myel | 10 (6) |
| l_transversetemporal_grayvol | 14 (0) |
| r_lingual_foldind | 15 (3) |
| l_rostralanteriorcingulate_meancurv | 15 (4) |
| r_parahippocampal_thck | 16 (10) |
| r_frontalpole_grayvol | 17 (7) |
| l_caudalanteriorcingulate_meancurv | 19 (4) |
| r_transversetemporal_area | 23 (21) |
| r_middletemporal_thck | 25 (15) |

Differently from the other factors, the direction of contribution of each feature in predicting Linguistic Skills is not in one major direction (cf. Figure 2d).

## 4 Conclusion

We aimed to predict individual differences in cognitive functioning only by using brain surface-based morphometry and cortical myelin estimates. Although this is a notoriously challenging problem, our regression model yielded good performance (Pearson-r correlation of almost 0.8) in predicting a sex-related factor, and almost a satisfactory performance (Pearson-r correlation around 0.11) in predicting fluid intelligence, visual memory and linguistic skills (cf. Table 3). These results highlight the potentiality of these type of features being used to predict cognitive performances.

We could not achieve fair performance or stability in predicting Executive Functions, Sustained Attention, Waiting Impulsivity, Verbal Episodic Memory and Visuo-spatial Skills, hinting that structural neuroimages alone cannot predict these cognitive factors, and other types of data are probably needed.

We consider that the approach described in this paper shows that the usage of specific, well-defined and self-explanatory features can help in the quick identification of meaningful regions of interest that are important to interpret machine learning methods, avoiding visual interpretation of neuroimages, which do not convey as much meaning as the features we extracted.

From our analysis it was possible to see that while some regions of interest appear more often, others do not appear so often thus suggesting that they might not be as important for understanding general cognition. As well, some specific features seem to not be important as they never appeared in the lists of most important features (eg. number of vertices, integrated rectified gaussian curvature, and intrinsic curvature index). It was also interesting that some factors had a clear difference of which hemisphere contributes more.

The significance of these results are supported in the fact that we used HCP, which is not only one of the most homogeneous and well characterised open datasets for healthy subjects, but also has an impressive number of people to be analysed (around a thousand), bringing more credibility to the results we presented.

We hope our work could serve as inspiration for researchers in the field to better study differences of cognition using well-implemented machine learning models that can be easily and correctly interpreted.

## Acknowledgements

Data were provided in part by the Human Connectome Project, WU-Minn Consortium (Principal Investigators: David Van Essen and Kamil Ugurbil; 1U54MH091657) funded by the 16 NIH Institutes and Centers that support the NIH Blueprint for Neuroscience Research; and by the McDonnell Center for Systems Neuroscience at Washington University. Tiago Azevedo is funded by the W. D. Armstrong Trust Fund, University of Cambridge, UK. Luca Passamonti is funded by the Medical Research Council grant (MR/P01271X/1) at the University of Cambridge, UK.

## Appendix

### Naming of brain features

The notation used in this paper to identify the brain-extracted feature consist of three parts separated by underscores. The first part is the hemisphere (**l** for left and **r** for right). The second part is short form of a region of the brain as defined by the Desikan-Killiany Atlas [2]. The third part is the specific feature extracted from that region of the brain, namely **thck** (cortical thickness), **area** (surface area), **numvert** (number of vertices in the cortex), **grayvol** (gray volume), **meancurv** (integrated rectified mean curvature), **gauscurv** (integrated rectified gaussian curvature), **foldind** (folding index), **curvind** (intrinsic curvature index), and **myel** (myelin estimate).

5 https://www.humanconnectome.org/study/hcp-young-adult/document/1200-subjects-data-release

6 http://surfer.nmr.mgh.harvard.edu/

7 https://www.humanconnectome.org/study/hcp-young-adult/article/s1200-group-average-data-release

